# Visualising uncertainty in brightfield to fluorescent image inference with *BFNet*

**DOI:** 10.1101/2020.03.13.987420

**Authors:** Scott Warchal, Célia Gautier, Thierry Dorval, Jean-Philippe Stephan

## Abstract

Predicting fluorescently labelled cellular structures from brightfield images is a recent application of convolutional neural networks and has already proven a valuable tool in biological imaging studies. These methods reduce the need for time-consuming manual annotations for supervised datasets, potentially reduce the costs in high-throughput imaging screens, and open up a number of potentially novel analyses. However as with any prediction there can be sources of error and uncertainty. Here we present BFNet, a method to visualise the uncertainty in predicted images by applying Monte Carlo dropout during inference and calculating per-pixel variance as an uncertainty map. Our method demonstrates the ability to highlight regions of an image where prediction is difficult or impossible due to imaging artefacts such as occlusions or out-of-focus images, as well as more general uncertainty when a trained model is applied to new data from different imaging of experimental settings. We have provided a python implementation of the method which is available at github.com/swarchal/bfnet.

## Main text

In 2018 two studies demonstrated that the information contained within brightfield images of cells can be used to infer specific subcellular structures by training U-Net^1^ inspired convolutional neural networks on paired datasets of brightfield images and their corresponding images with fluorescently labelled cellular structures^2,3^. The overall performance of these models can be assessed during training by calculating the mean-squared-error, or other appropriate loss functions, between the real and predicted fluorescent images on a withheld validation dataset, and after training on a separate test dataset. Whilst this gives a global measure of how well a model is performing on a labelled dataset, it does not give researchers any information on how well they can trust the individual predictions on new unlabelled data, and perhaps more interestingly – how well they can trust local regions within a predicted image.

Our method is based on prediction stability, i.e. if a prediction is robust then small changes to model weights or the input itself should not drastically alter the output, therefore regions in the predicted output image that are highly variable in the presence of small perturbations are less trustworthy. This is inspired by the work of Gal *et al*. who demonstrated that Bayesian neural networks, which accurately capture model uncertainty but are difficult to implement and train, can be approximated by Monte Carlo dropout applied to the weights in a neural network during inference^4,5^. Prediction uncertainty is obtained through multiple stochastic forward passes for each input, and the variance is calculated on the collected predictions. This approach to generate uncertainty maps for predictions has recently been used by Soleimany *et al.* for the task of parasite segmentation in images of liver stage malaria^6^, but has not yet been applied to brightfield to fluorescent image inference.

**Figure 1:**
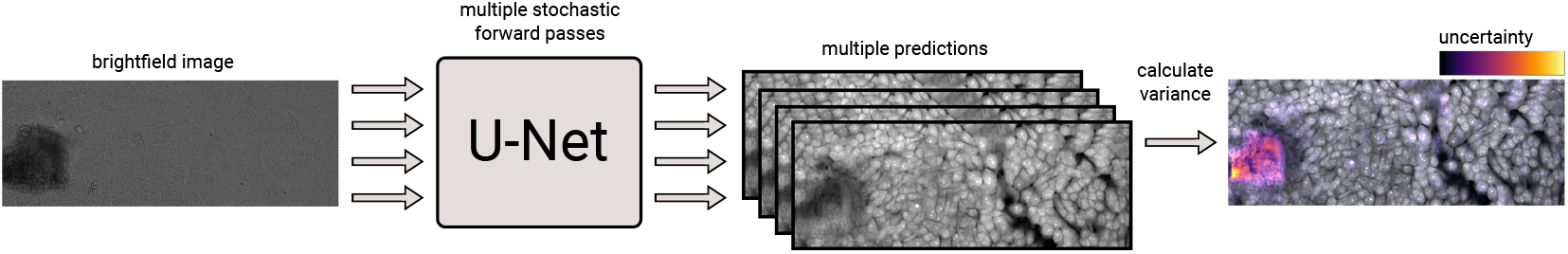
A diagram of brightfield to fluorescence image inference with overlaid uncertainty map. A single brightfield input image is passed as input to a trained U-Net model where it is subjected to multiple forward passes with dropout (either weight or pixel dropout), after which the pixel-wise variance is calculated from the multiple predictions and visualised as an uncertainty map when overlaid on the predicted fluorescence image.

We released our method as a python package using the scikit-image, numpy and pytorch packages. It contains two different methods of calculating the uncertainty map: (i) the first method uses weight dropout during inference, whereby dropout layers are present in all convolutional blocks of the U-Net model. Entire channels are randomly zeroed independently at each forward pass with a probability sampled from a Bernoulli distribution^7^. After multiple forward passes for a single input image, the pixel-wise variance is calculated from the collected predictions and used to generate an uncertainty map. The default behaviour of pytorch is to adjust the function of certain layers (dropout, batchnorm etc.) depending on whether the model is in training or inference mode, therefore we have to alter the behaviour of the dropout layers to enable dropout during inference and disable it during training without altering the behaviour of the other layers. (ii) The second method works in a similar way, but applies the dropout to the pixels in the input image rather than the network weights, by zeroing a small proportion of pixels in the image after image normalisation followed by multiple forward passes and a variance on the collected predictions. We found this method of dropout performed on the input images more sensitive to the chosen hyper-parameters, although it has a benefit of being agnostic to the neural network library and can be easily wrapped around existing trained models without any modifications to the underlying network architecture.

To obtain a prediction there is a choice between predicting using no dropout, or by taking an average of the stochastic forward passes with dropout. In our experience, we found both methods work satisfactorily with largely similar results, although the averaged prediction tends to appear more blurred. We also observed that the uncertainty maps produced by either method highlight regions in the brightfield image that are ambiguous due to dust occlusions or cells growing in a non-monolayer. When we subjected the brightfield images to artificial perturbations such as black/white squares or colour inversions, we found that the uncertainty maps typically highlight these regions where prediction is difficult; although the method performs poorly with local Gaussian blurring or instances where an occlusion covers a large portion of the image (see supplementary data).

Here, we present a method to quantify and visualise uncertainty when predicting fluorescent labels from brightfield images. As brightfield to fluorescent image inference has potential applications in high-throughput imaging studies, we hope that the community can use this method further reason about their predictions, as well as enabling researchers to subject potentially troublesome predictions to further scrutiny. Our method is released as BFNet, an open source python package which the community is encouraged to use and adapt to their needs.

## Supplementary data

### Typical workflow

Here we show a trained BFNet model used to predict Hoechst from an unlabelled brightfield image. The variance map can be inspected as a separate channel or as an overlay on the brightfield image to show regions in the input where the model is uncertain, or on the prediction to show where the predictions should be further scrutinised.

**Supplementary Figure 1:**
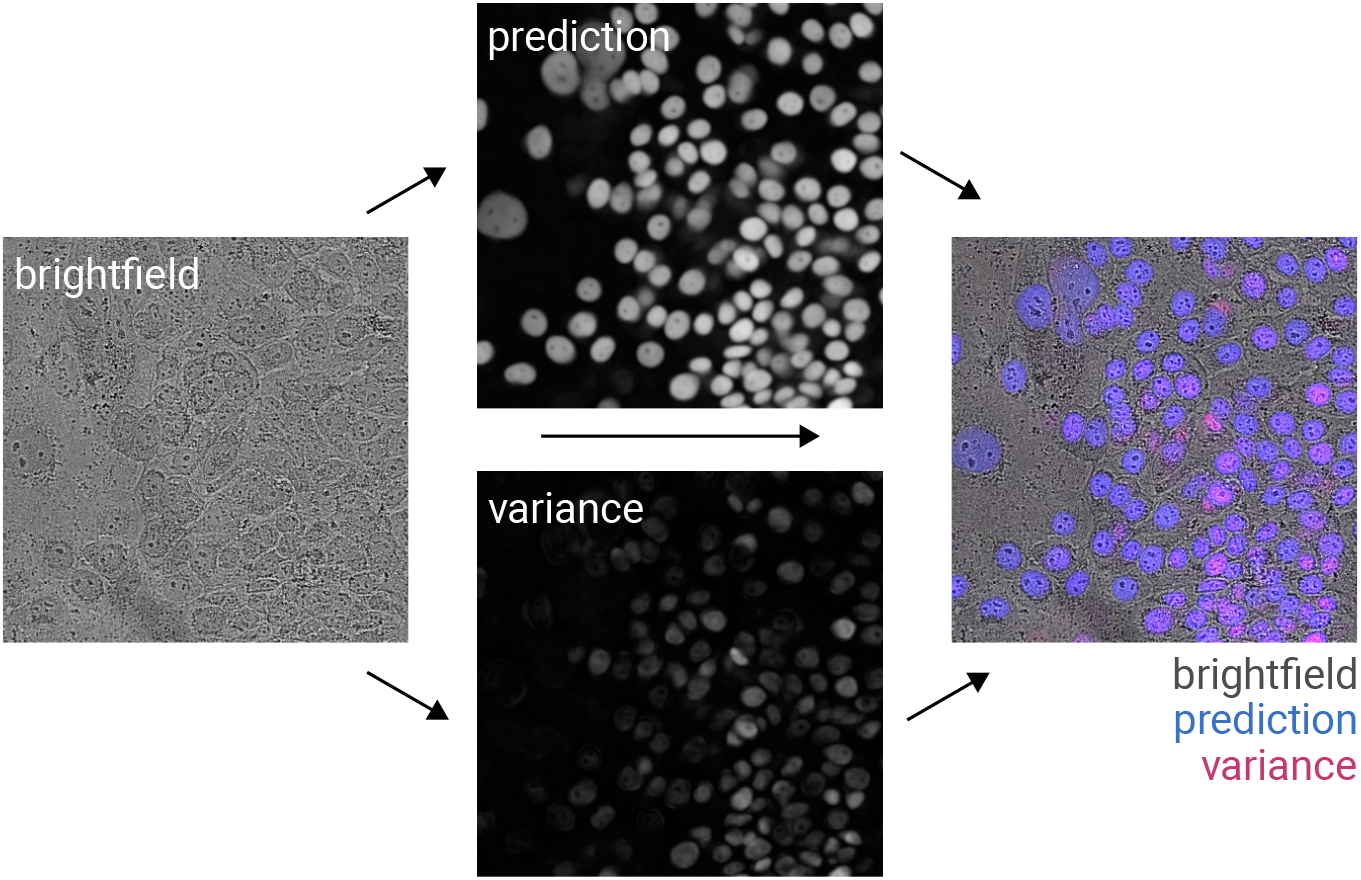
Typical workflow showing the prediction of Hoechst staining from an unlabelled brightfield image along with the calculated variance used as an uncertainty map. This can then be overlaid over the prediction or the brightfield image or both. The variance map was calculated using weight dropout with a proportion of 0.1 dropout and 20 stochastic forward passes. The prediction image shows the mean of those 20 forward passes. The brightfield images in this figure have been subjected to localized contrast enhancement (CLAHE) in ImageJ for visualisation purposes.

### Performance on artificially perturbed images

To determine the performance of the method, several brightfield images were altered in ImageJ to introduce various artefacts such as the addition of black or white areas, local or global blurring and colour inversion. Two models (pixel dropout and weight dropout) were then trained on the same dataset, which consisted of brightfield images paired with Hoechst and cellmask fluorescent staining, excluding the original images from which the altered images were created from. We can see that some of the perturbations do not result in regions of uncertainty despite areas of missing data and nonsense predictions, this is particularly evident for the images with localised blurring, or extremely large occlusions. Fortunately, perturbations which are more likely encountered in real microscopy environments such as global blurring to simulate an out-of-focus image and and small occlusions result in expected regions of high variance signalling uncertainty in the predictions.

**Supplementary Figure 2:**
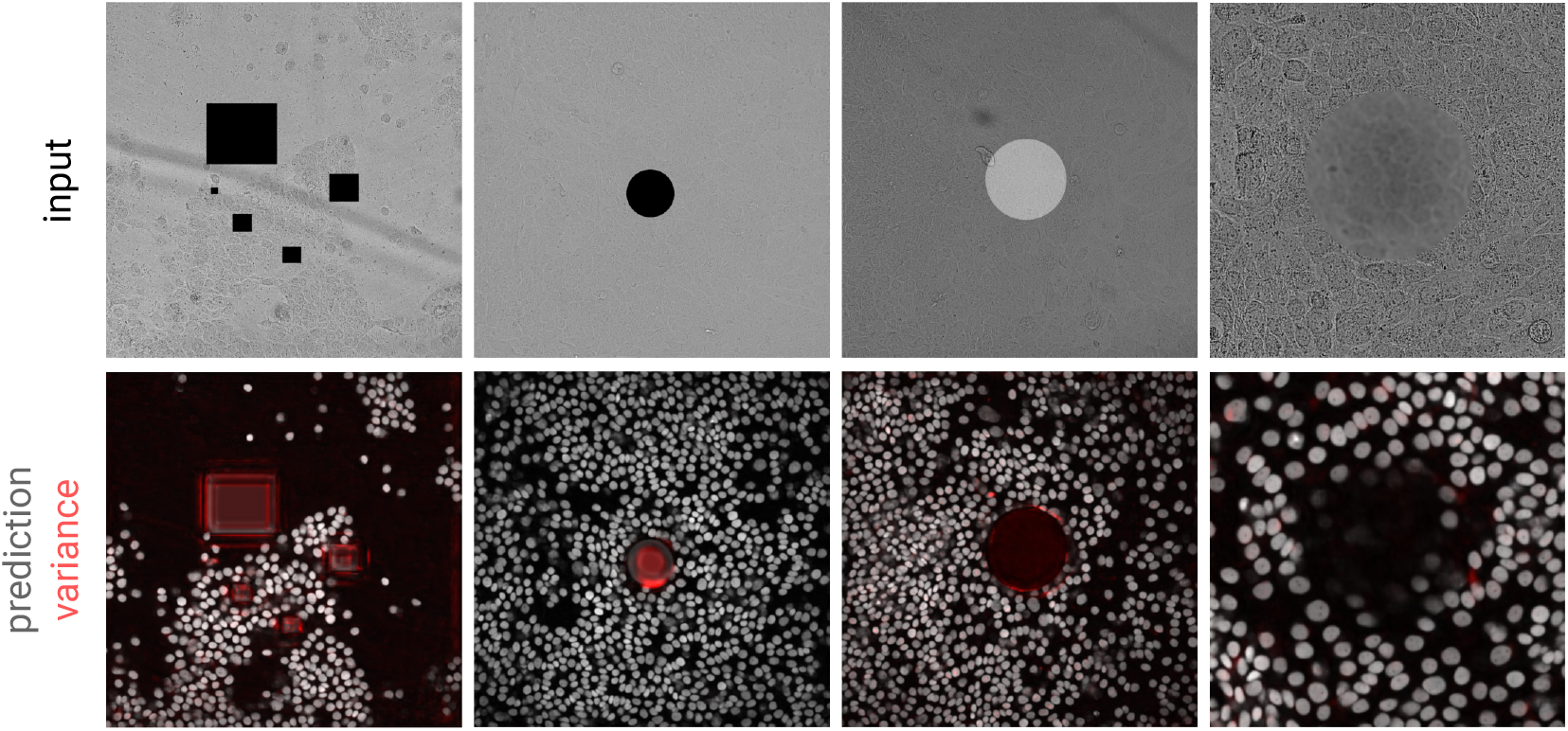
Performance on artificially perturbed images. Hoechst predictions from brightfield images modified to introduce regions of missing or out-of-distribution data. Predictions are shown below with uncertainty overlaid in red.

### Correlation between prediction intensity and variance

It is a concern that variance appears to be highly correlated with the intensity of the predicted pixels. We have plotted pixel-intensity of the prediction against the variance of that pixel (Supp. fig. 3). We see that there is indeed some correlation between pixel intensity and the variance value. We think this because pixel intensity in fluorescent images typically have a log-normal distribution, and so bright objects have a much greater range in which their values can vary compared to dim objects. It may be possible to alleviate this problem through scaling the images during the variance calculation, but this has yet to be investigated properly.

**Supplementary Figure 3:**
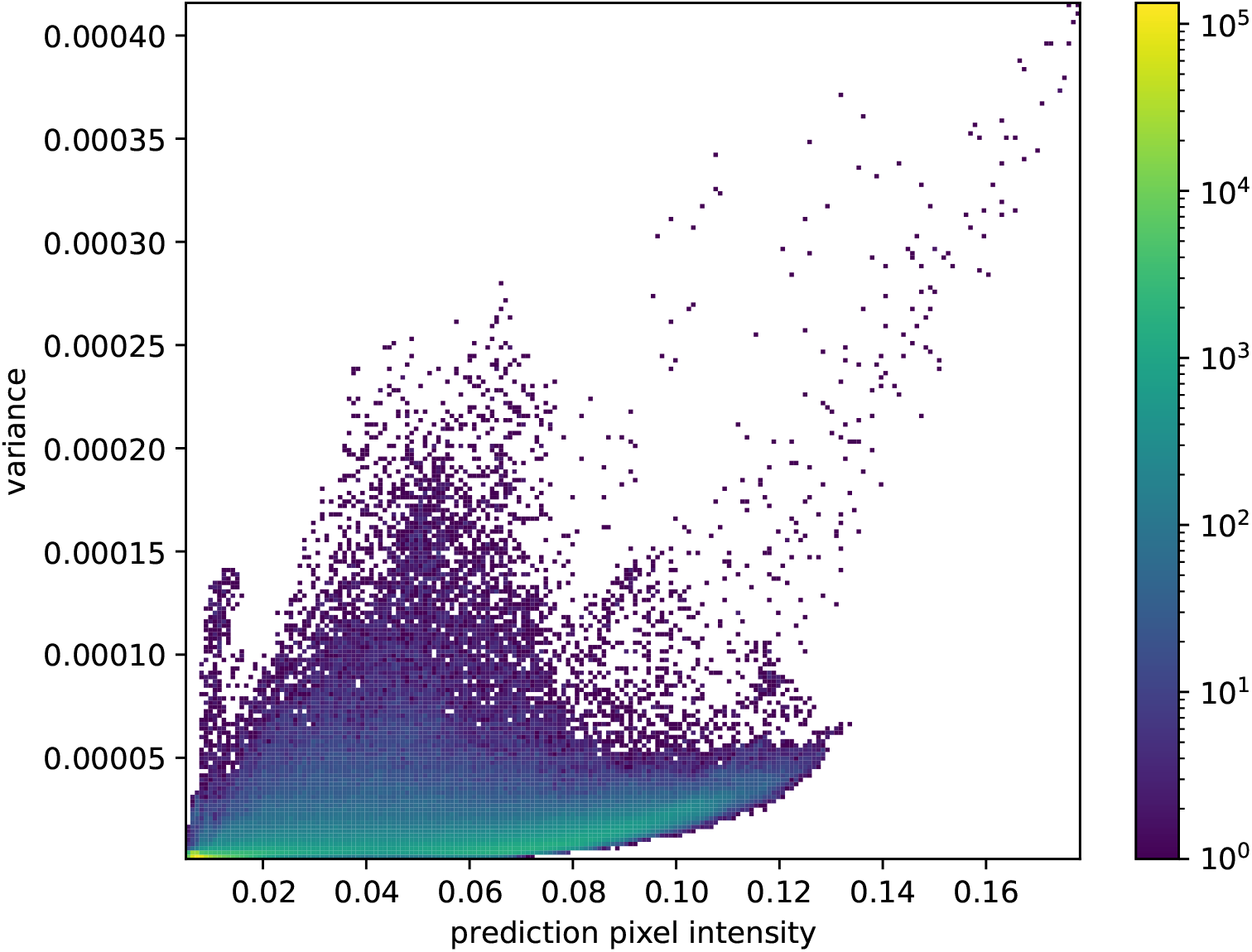
2D histogram of prediction intensity plotted against variance per pixel for a single Hoechst prediction.

### Comparison between predictions from a dropout average versus no dropout

Since the variance calculation uses multiple forward passes, it is possible to calculate the predicted image from an average of these predictions. We that the average images from multiple predictions with dropout have no benefit over more typical predictions using a single forward pass with no dropout.

**Supplementary Figure 4:**
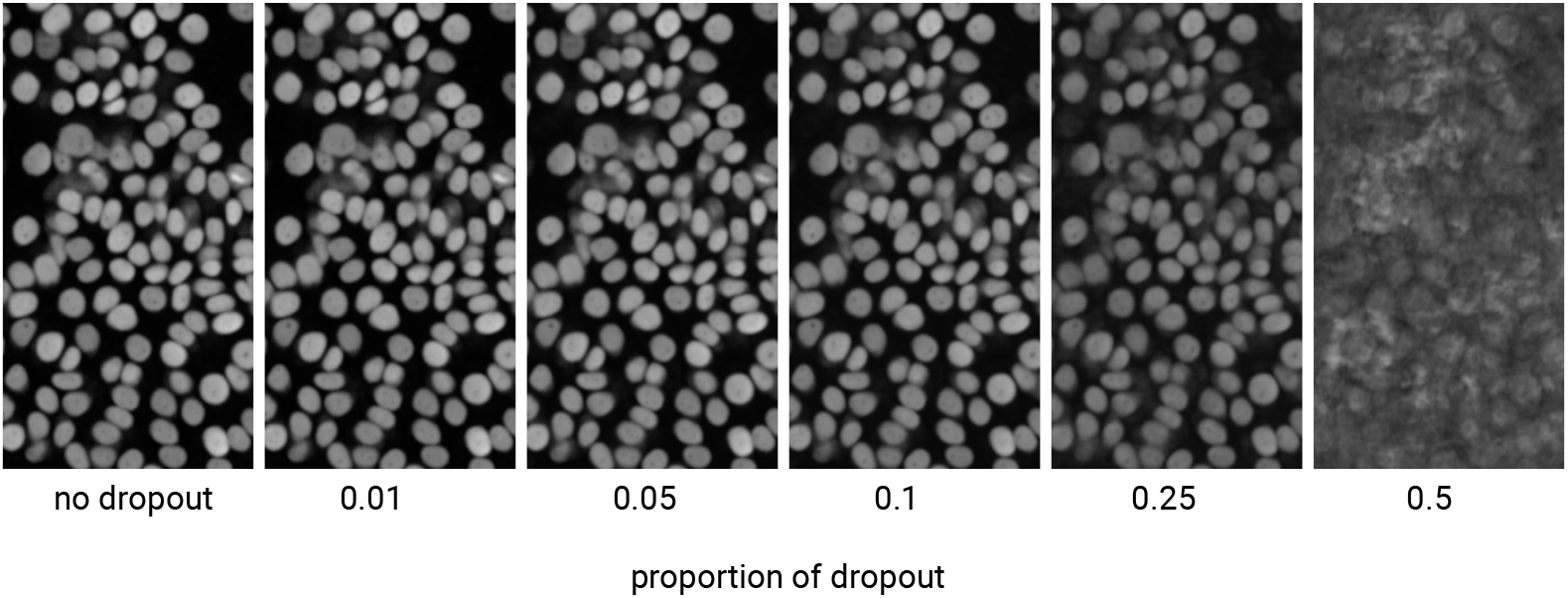
Comparison between predictions from an average of multiple forward passes versus a single forward pass with no dropout. The left-most image shows the predicted Hoechst image from a single forward pass with no dropout. The subsequent images then show the mean of 20 forward passes with differing proportions of dropout indicated below each image.

### Hyperparameters

#### Weight dropout

The weight dropout method has two hyperparameters which can be adjusted at inference time: the proportion of dropout, and the number of stochastic forward passes. The default for these values are a dropout proportion of 0.1 and number of forward passes of 20. Increasing the number of forward passes generally increases the accuracy of the variance calculation although this linearly increases the computation time. The model is quite sensitive to the proportion of dropout, with larger values (≥ 0.25) producing a more coarse uncertainty map, and lower values (≤ 0.1) highlighting smaller regions typically sub-cellular in size (Supp. fig. 5).

**Supplementary Figure 5:**
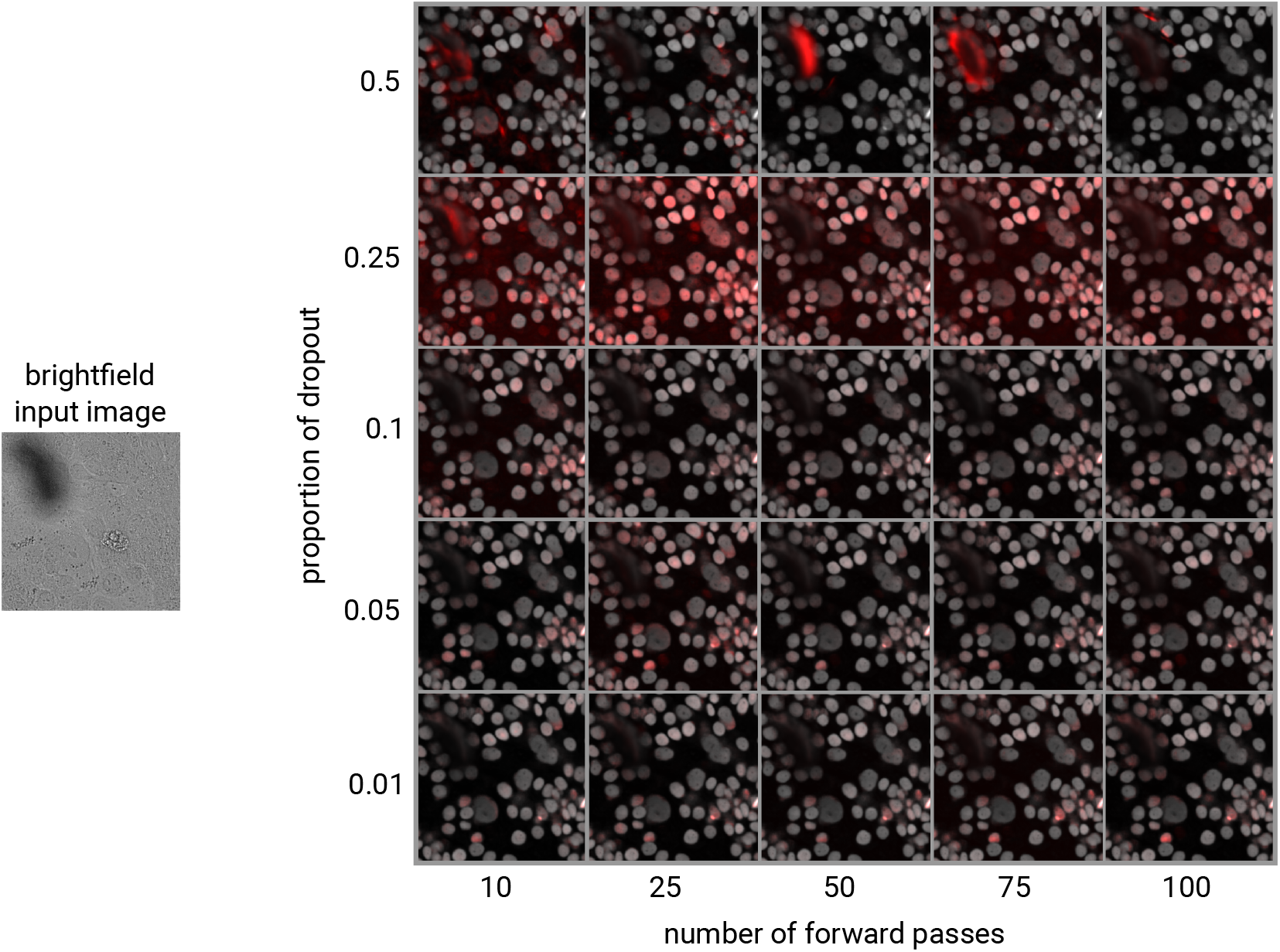
Parameter grid showing the influence of adjusting the proportion of dropout and the number of forward passes on the calculated variance shown in red, and predicted Hoechst staining shown in grayscale. The same trained model was used to each prediction, and predictions were made from a single forward pass with zero dropout.

